# Reproductive success in the barn owl is linked to partner compatibility in glucocorticoid levels

**DOI:** 10.1101/517227

**Authors:** Paul Béziers, Lukas Jenni, Alexandre Roulin, Bettina Almasi

## Abstract

In biparental species, reproductive success depends not only on the quality of the parents, the care they each provide and many environmental factors such as territory quality and food availability, but also on the ability of the parents to collaborate and divide reproductive tasks. Because hormones, such as corticosterone (CORT), modulate physiological and behavioural functions that are associated to reproductive success, hormonal compatibility between pair members is likely to have consequences on reproductive success. Here, we investigated in the barn owl (*Tyto alba*) whether baseline and stress-induced CORT levels are correlated between breeding partners and whether this correlation is associated to fitness parameters (*i.e*., clutch size, offspring number and body mass). We found that the combination of CORT levels in the two partners predicts reproductive parameters. Pairs with similar baseline CORT levels during incubation produced more fledglings than pairs with dissimilar baseline CORT levels. On the other hand, pairs showing dissimilar stress-induced CORT responses during the period of offspring rearing produced more fledglings than pairs with similar stress-induced responses. Offspring body mass was associated only with maternal (baseline and stress-induced) CORT levels and depended on the context with baseline CORT being potentially adjusted to environmental conditions but also to the energetic demand of reproduction. Hence, to produce large broods good quality mothers might increase their baseline CORT to meet the energetic demand of the brood, while mothers in unfavourable habitats may have smaller broods but still need to increase baseline CORT to cope with the environmental challenges. Taken together, the results show that the association between parental CORT levels and reproductive success are context-dependent and rely on the combination of parental CORT levels. Assuming that CORT levels reflect investment in parental care, our study suggests that parents coordinate their reproductive activities in a complex way to ensure a high reproductive success.

## Introduction

Reproductive success can vary greatly between individuals and populations as it depends on many intrinsic and extrinsic factors. In species with biparental care, reproductive success depends on the genetic and phenotypic quality of both partners and on how reproductive tasks are divided among them. However, a partnership is a dynamic interaction driven by an agreement over how partners divide their reproductive investment. This depends on the level of compatibility and coordination between partners (Ihle et al. 2015) and how cooperative they are and negotiate their relative investment in reproduction (McNamara et al. 1999). For instance, partners can benefit if they complement each other in the sharing of reproductive tasks, such as incubation, nest defense and offspring provisioning (Both et al. 2005; Burtka and Grindstaff 2015; Dingemanse et al. 2004; Fox and Millam 2014; Gabriel and Black 2012; Itzkowitz et al. 2002; Laubu et al. 2016; Schuett et al. 2011a; Spoon et al. 2006). Dissimilarity in behaviour or personality between partners can also be an advantage in species where partners have a different role in parental care, because it can reduce intra-pair competition and increase the efficiency of parental tasks, for example, in how the territory is exploited (Wolf and Weissing, 2010, 2012). For instance, in cockatiels (*Nymphicus hollandicus*) male and female partners with mixed personalities are less aggressive to each other and better coordinate incubation activities, which ultimately leads to a higher hatching success (Fox and Millam 2014). Although partner compatibility or dissimilarity seems to be an important factor of reproductive success, the underlying mechanism is usually unknown.

Social and sexual behaviours are often regulated by hormones (Becker et al. 2002; Oliveira 2009) and hence to coordinate reproductive activities, partners should coordinate the regulation of their hormone profiles (Hirschenhauser 2012). For instance, in greylag geese (*Anser anser*) and great tits (*Parus major*), partners showing similar testosterone and corticosterone profiles (and personalities in tits) during nesting care and throughout the year are less likely to divorce and achieve a higher reproductive success (Both et al. 2005; Hirschenhauser et al. 1999; Ouyang et al. 2014; Weiss et al. 2010). Inversely, in convict cichlids, *Amatitlania siquid*, partners with dissimilar glucocorticoid (cortisol) levels produced more eggs than partners with similar glucocorticoid levels (Schweitzer et al. 2017).

Glucocorticoid hormones, such as corticosterone (CORT), play an essential role in the maintenance of body functions by modulating energy expenditure and behaviour in response to environmental challenges (Breuner et al. 1999). Baseline CORT levels follow daily and seasonal rhythms that match the different needs and behaviour of the life cycle routine. In response to unpredictable perturbations in the environment, CORT levels can rapidly increase to redirect resources towards essential functions, such as locomotion and foraging activity (e.g., Almasi et al. 2008; Angelier et al. 2008; Bonier et al. 2011; Crossin et al. 2012), which consequently affect reproductive success (see Bonier et al. 2009a; Breuner et al. 2008 for review). Because CORT regulates many physiological, behavioural and reproductive functions, it is often associated with personality traits, and hence may partly mediate how behaviours of breeding partners are coordinated (e.g. Both et al. 2005; Burtka and Grindstaff 2015; Fox and Millam 2014; Gabriel and Black 2012; Laubu et al. 2016). This is so, because CORT can regulate parental behaviour and help respond to reproductive challenges. However, the relationship between CORT and fitness components can be context-dependent, and therefore may vary between populations (Doody et al. 2008; Müller et al. 2007), years (Chastel et al. 2005; Henderson et al. 2017; Lanctot et al. 2003), sexes (Angelier et al. 2010) and reproductive strategies (Lancaster et al. 2008) or depend on the life-history stage at which individuals have been sampled (Bonier et al. 2009a; Ouyang et al. 2011).

Increasingly, it is recognized that behavioural and hormonal compatibility between partners plays a role in reproductive success (Both et al. 2005; Burtka and Grindstaff 2015; Gabriel and Black 2012; Hirschenhauser 2012; Ouyang et al. 2014; Schuett et al. 2011a; Schweitzer et al. 2017; Spoon et al. 2006). Given that CORT is essential in the regulation of behaviours associated to reproduction (Love et al. 2014), including incubation behaviour (Schoenle et al. 2017; Thierry et al. 2013) and offspring feeding rate (Almasi et al. 2008; Ouyang et al. 2013a), we investigated the similarity of baseline and stress-induced CORT levels between the breeding partners of barn owls (Tyto alba) and how similarity or dissimilarity is related to reproductive success. In this medium-sized bird, only the female incubates the eggs and the male forages alone for the family until the offspring are approximately two weeks of age when the female starts to participate in foraging. Previous experimental studies have shown that high CORT levels decrease feeding rates of male and female barn owls (Almasi et al. 2013; Almasi et al. 2008) and is associated with survival of adults (submitted). Here, we examine whether baseline CORT and stress-induced CORT response of pair members are associated with reproductive parameters including clutch size, brood size at fledging and nestling body mass.

## Materials and methods

### Study species

From 2004 to 2016, we studied a population of barn owls in western Switzerland (46°46′42″ N, 6°38′28″ E) in an area of 1070 km^2^ dominated by an agriculture landscape at an altitude of 420 to 730 m. The barn owl is a medium-sized raptor (mass of adults 239 to 515 g) which preys on small mammals and produces one to two annual clutches per year between February and August (Béziers and Roulin 2016). Females lay 2 to 12 eggs per clutch (mean 6.3 ± SD 1.5) every 2-3 days and females start incubating once the first egg has been laid, which leads to a pronounced hatching asynchrony. Females incubate the eggs and stay in the nest cavity to take care of the offspring until they are able to thermoregulate by themselves. Therefore, males hunt alone until the oldest nestling is between 2 to 3 weeks old when the mother starts to participate in foraging for the progeny.

### Study area

Since 1990 we have fixed 360 nest-boxes in barns and recorded laying date, clutch size, number of hatchlings and brood size at fledging. Adult barn owls (118 males and 121 females) were captured during the day or at night while the female was incubating the eggs or the parents were provisioning the offspring. For each adult, we measured body mass (to the nearest g) and wing length (to the nearest mm) and took a blood sample for hormonal and molecular analyses. Because only females incubate the eggs and hence have a brood patch, we identified the sex of adults by the presence or absence of a brood patch. The age of 103 males and 108 females was known because they were ringed as nestlings, while for 15 males and 13 females ringed as adult we estimated their age from their moult pattern (Taylor 1993). For the statistical analyses, we classified birds as “yearlings” (*i.e*., individuals in their first-year of life) or “adults” (*i.e*., individuals older than one year).

During the nest visits, we ringed all nestlings (299 males and 315 females from 143 different broods) and measured their body mass and wing length. Hatching date was estimated by using wing length (Roulin 2004). A blood sample was taken to determine nestling sex (Roulin et al. 1999). To disentangle genetic from environmental factors on CORT levels, we cross-fostered eggs or nestlings between pairs with similar laying dates. If clutch size between pairs of nests was of similar size (± 1 egg), we swapped all eggs, whereas if the difference was larger than one egg or if the difference in laying date between cross-fostered pairs was too large (*e.g*., 12 days), we either cross-fostered the same number of eggs at the same stage of incubation or swapped the oldest nestlings from one nest with the youngest nestlings of the other nest. In any case, we kept intact the within brood age hierarchy and brood size (± 1 egg).

### Blood sampling and hormone analyses

To determine corticosterone levels of adult barn owls, we used a standard capture-restrain protocol (Wingfield et al. 1982), *i.e*. birds were captured and a first blood sample was quickly taken to measure baseline CORT levels and later a second blood sample was collected to measure stress-induced CORT levels. Adult females were sampled between 8:16 in the morning and 24:00 in the evening, while males were sampled over 24 hours. The blood was taken by puncturing the brachial vein and collected with heparinised capillary tubes. After bleeding, the plasma was directly separated from blood cells and flashed frozen in liquid nitrogen. Once back from the field within 24 hours the samples were stored at −20°C until analysis. The first sample was taken between 30″ and 3′19″ (1′25″ ± 34″5 average sampling time) after capture and was considered to represent the baseline level of corticosterone (93 breeding male and 93 breeding female samples). The time laps between capture and blood sampling had a marginal effect on baseline corticosterone levels (*β* = 0.13, 95% credible interval (95% CrI) [0.02; 0.23], Table S1). The individuals were then held in an opaque cloth bag and bled again approximately 25 minutes after capture (24′58″ ± SD 1′56″) to measure the corticosterone response to handling stress (116 male and 116 female samples). This time lapse represents the peak of corticosterone response in the barn owl (Almasi and Roulin 2015) and was not associated to stress-induced CORT levels (*β* = 0.15 95% CrI [−4.07; 4.5], Table S1).

Plasma corticosterone was extracted with dichloromethane and determined with an enzyme immunoassay (Munro and Stabenfeldt 1984; Munro and Lasley 1988) following (Müller et al. 2006). Ten microliters of plasma were added to 190μl water and extracted with 4ml dichloromethane. The solution was mixed for 30 minutes on a vortex machine and then incubated for 2 hours. After separating the water phase from the dichloromethane, the dichloromethane was evaporated at 48°C and corticosterone was re-suspended in 550 μl of phosphate buffer. The dilution of the corticosterone antibody (Chemicon; cross-reactivity: 11-dehydrocorticosterone 0.35%, progesterone 0.004%, 18-OH-DOC 0.01% cortisol 0.12%, 28-OH-B 0.02% and aldosterone 0.06%) was 1:8000. We used HRP (1:400 000) linked to corticosterone as enzyme label and ABTS as substrate. We determined the concentration of corticosterone in triplicates by using a standard curve run in duplicates on each plate. If the corticosterone concentration was below the detection threshold of 1ng/ml the analysis was repeated with 15μl or 20μl plasma. If the concentration was still below the detection limit (1 baseline sample), we assigned to the sample the value of the assay detection limit (1ng/ml). Plasma pools from chicken with a low and high corticosterone concentration were included as internal controls on each plate. Intra-assay variation ranges from 5 % to 14 % and inter-assay variation from 7 % to 23 %, depending on the concentration of the internal control and the year of analysis.

### Statistical analyses

We analysed the relationship between CORT levels (baseline and stress-induced) and two different reproductive parameters (*i.e*., clutch size and number of fledglings). In our analysis, the number of successfully fledged offspring was the number of offspring having reached the age of fledging (55 days). These two variables were fitted as dependent variables in separate linear mixed effect models with the following covariates: parental CORT levels (baseline and stress-induced levels in separate models), the breeding stage at which individuals were sampled (late incubation stage or while provisioning the offspring (hereafter nestling care)), the age of parents (yearling vs. adult), the condition of parents (*i.e*., mass of the individuals divided by wing length) and laying date in Julian days and its quadratic term (*i.e*., laying date^2^). For all models, we included the three-way interaction between CORT levels of male and female partners with the breeding stage to test whether the relationship with reproductive success depended on the similarity of hormonal levels between partners, and whether this relationship changed according to breeding stage. All the covariates were standardized to compare their relative effects. The year, brood identity, identity of females and males (*i.e*., some individuals were caught at different broods and with different partners) were added as random terms in the models and removed if they did not explain any variance. Baseline and stress-induced CORT levels are, to some extent, correlated to each other (*r* = 0.25, 95% CrI [0.12; 0.38]), which could confound the interpretation of our results. The relation between baseline and stress-induced CORT did not depend on the breeding stage neither the sex of individuals (correlation estimates and 95% CrI within breeding stage and sexes were above 0). We also tested whether CORT levels and body condition (*i.e*., body mass/wing length) of partners were correlated and whether it differed according to breeding stage. Male and female CORT levels and body condition were standardized and correlation tests were done with *bayes.cor.test* function from *BayesianFirstAid* package (Bååth 2014). We also investigated the relationship between CORT levels of parents and body mass of nestlings aged 5 to 30 days. In this model, we considered the hour of sampling and its quadratic term (hour of sampling^2^), the Julian date of sampling, nestling age and sex and their position in the within-brood age hierarchy, brood size and breeding stage. We also included the maternal and paternal CORT levels (baseline and stress-induced levels in separate models), the three-way interaction between parental CORT levels with the breeding stage and the interaction of mother and father CORT levels with brood size (Table 2). The different models were assessed by simulating 50’000 random values from the joint posterior distribution of the model parameters. From these simulated data, we used the 2.5 % and 97.5 % quantiles as the lower and upper bands of our 95% credible interval (CrI) (Korner-Nievergelt et al. 2016). We visually inspected each model to check whether they respected the model assumptions (normality, homogeneity). For the figures, the regression lines and 95% CrI were drawn from 5’000 simulations of pairs of intercepts and slopes based on the posterior distribution of the model parameters. All models and figures were performed in R version 3.4.3 (R Core Team 2018) with the *lmer4* (Bates et al. 2015) and *arm* packages (Gelman and Hill 2007). The analyses for baseline and stress-induced CORT levels did not include necessarily the same individuals as we were not always able to obtain both samples from all individuals. Thus, over the 152 and 187 individuals used in the baseline and stress-induced CORT analyses, respectively, only for 100 of them could we measure both baseline and stress-induced CORT levels.

## Results

### Corticosterone levels and body condition of breeding partners

Baseline CORT levels between pair members were positively correlated (Figure 1, correlation during incubation (*r* = 0.28, 95% CrI [−0.02; 0.53], posterior probability that the correlation between partner was positive 96%) and nestling care (*r* = 0.54, 95% CrI [0.30; 0.73]); a similar conclusion applies to stress-induced CORT levels between pair members (during incubation (*r* = 0.47, 95% CrI [0.19; 0.67]) and nestling care (*r* = 0.26, 95% CrI [0.02; 0.48]). On the other hand, body condition of male and female partners was positively associated during nestling care (Figure 2; *r* = 0.24, 95% CrI [−0.01; 0.50], posterior probability that the correlation between partner was positive 94%) but not during incubation stage (*r* = −0.01, 95% CrI [−0.34; 0.16]).

**Figure 1.**
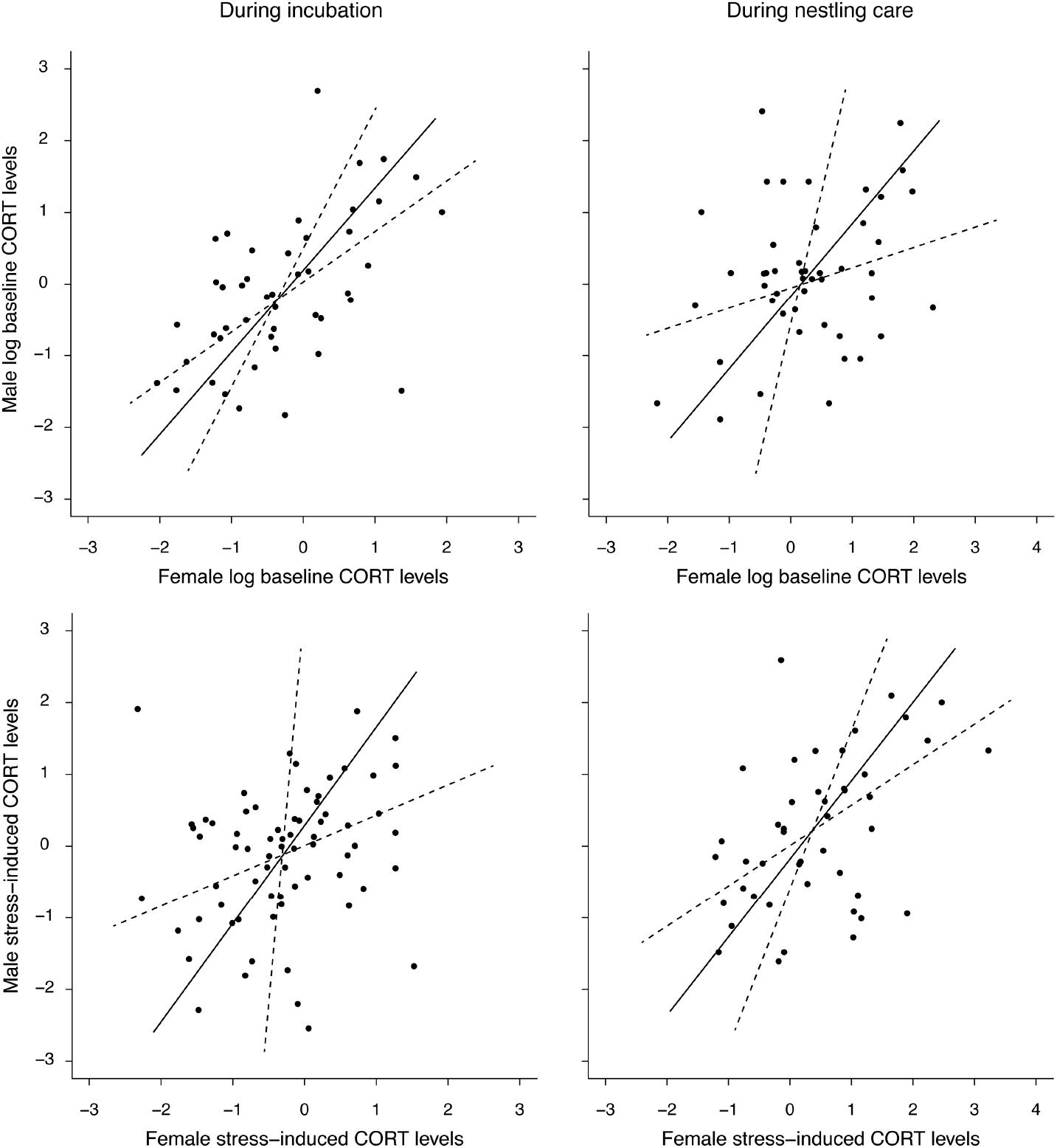
Baseline and stress-induced CORT correlation for members within breeding stage. Baseline and stress-induced levels of both males and females were standardized. The black lines represent the major correlation axes and the dotted lines the 95% confidence intervals.

**Figure 2.**
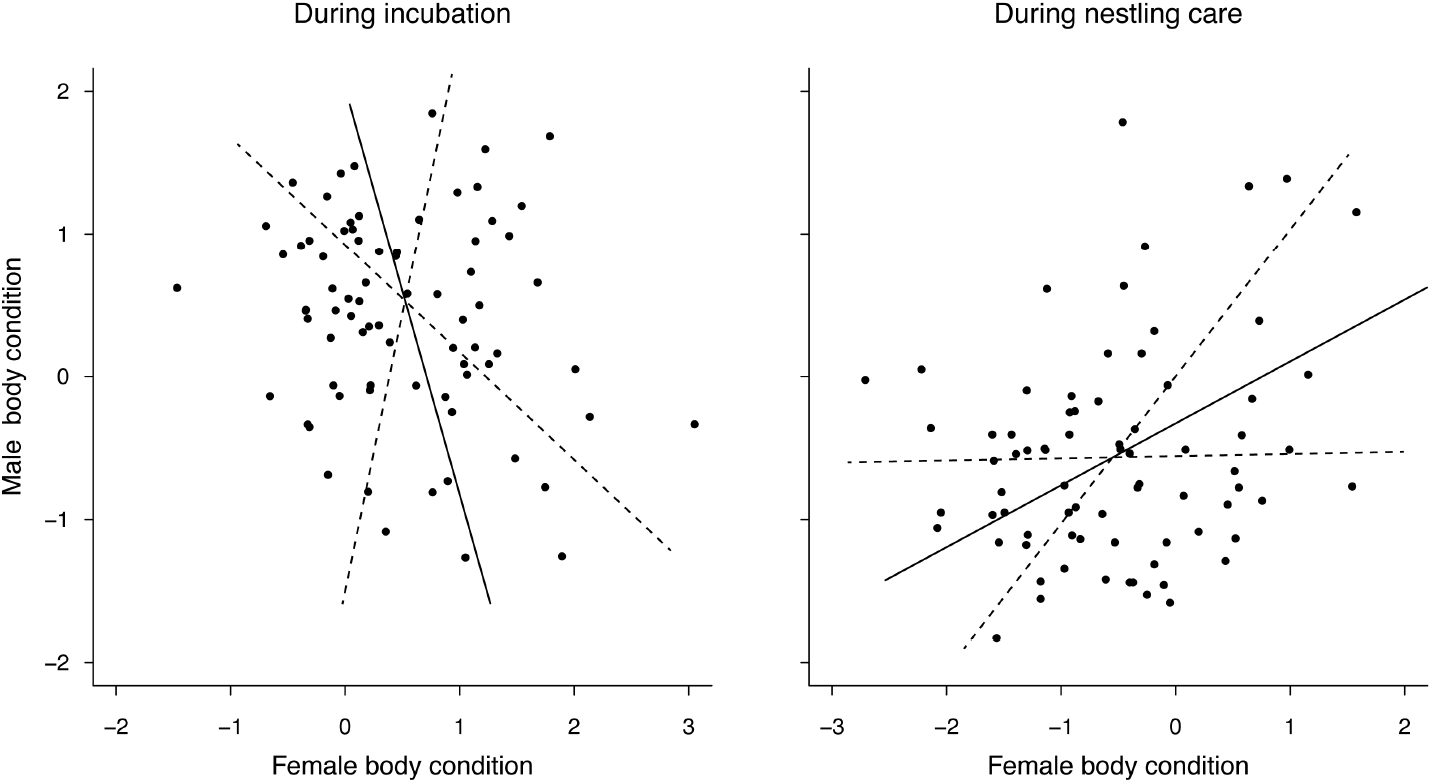
Body condition (mass/wing) correlation between pair members within breeding stage. The black and the dotted lines represent the mean posterior distribution estimated from the model for during incubation and nestling provisioning, respectively. The black lines represent the major correlation axes and the dotted lines the 95% confidence intervals.

### Corticosterone levels in relation to reproductive parameters

Clutch size did not depend on the similarity or dissimilarity of CORT levels of partners but was associated to stress-induced CORT of males in interaction with breeding stage (Table 1a). Males with lower stress-induced CORT response during incubation stage had larger clutches than males with a higher stress-induced response (Figure 4). A relation that did not prevail with the stress-induced CORT response measured during nestling care (*β* = 0.25, 95% *CrI* [−0.25; 0.73]). Number of fledglings was related to baseline and stress-induced CORT depending on the breeding stage at which adults were sampled (Table 1b). Partners with similar baseline CORT (both high or both low levels) during the late incubation stage produced more fledglings than partners with dissimilar baseline CORT, an effect that was not observed for baseline CORT sampled during nestling care (Figure 3). In contrast to baseline, stress-induced CORT in females during incubation was not related to the number of fledglings, whereas males with lower stress-induced CORT produced more fledglings than individuals with a higher stress-induced CORT response. During nestling care partners with similar stress-induced CORT levels had a lower number of fledglings than partners with dissimilar stress-induced CORT levels (Figure 5), an effect that was even stronger for females paired with males with very high CORT levels (figure not shown).

**Table 1.** Baseline and stress-induced CORT levels in relation to (a) clutch size and (b) number of fledglings of barn owl breeding pairs. Effects of different parameters are estimated from a linear mixed effect model based on 93 and 115 different breeding pairs measured during 11 and 9 years between 2004 and 2016 for baseline and stress-induced CORT levels, respectively. Estimates (95% Bayesian credible intervals) of predictors are based on the draw of 50’000 random values from the joint posterior distribution of the model parameters. The year (Tables 1), female (Tables 1a & 1b) and male identity (Table 1b) were added in the models as random terms to correct for pseudoreplication. Predictors with a significant effect on fitness components are written in bold. Adult females laid fewer eggs than yearling females, while in males the opposite pattern was detected. Clutch size decreased from the beginning to middle of the breeding season and increased again at the end of the breeding season. Although, the intervals of confidence for age and laying date were larger in the stress-induced models, their association with clutch size followed the same direction as in the baseline CORT model. Males in good condition (*i.e*., mass/wing) had also larger clutches than males in poorer condition; however, this effect was only observed in the baseline model (Table 1a). Females captured in poorer condition were, on the other hand, found to produce more fledglings than females captured in good condition, an effect that was clearly observed in the baseline model while in the stress-induced CORT model it was only a trend (Table 1b).

|  | *Model with baseline CORT* | *Model with stress-induced CORT* |
| --- | --- | --- |
| Fixed effects | *estimates<sup>†</sup> (95% CrI)* | *estimates<sup>†</sup> (95% CrI)* |
| Intercept | 6.17 (5.23 – 7.09) | 6.35 (5.56 – 7.14) |
| Male CORT | 0.21 (-0.31 – 0.73) | **-0.48 (-0.86 – -0.09)** |
| Female CORT | -0.03 (-0.58 – 0.52) | -0.13 (-0.54 – 0.28) |
| Breeding stage* | -0.02 (-0.96 – 0.93) | 0.06 (-0.66 – 0.78) |
| Laying date | -3.04 (-5.59 – -0.51) | -1.55 (-3.47 – 0.38) |
| Laying date<sup>2</sup> | 3.53 (1.16 – 5.903) | 1.66 (-0.26 – 3.591) |
| Female condition | -0.16 (-0.54 – 0.22) | 0.23 (-0.1 – 0.55) |
| Male condition | **0.40 (0.02 – 0.79)** | -0.04 (-0.37 – 0.29) |
| Female age class** | **-0.89 (-1.64 – -0.13)** | -0.52 (-1.15 – 0.09) |
| Male age class** | **1.20 (0.31 – 2.1)** | 0.17 (-0.41 – 0.76) |
| Male CORT x Female CORT | 0.01 (-0.46 – 0.47) | -0.01 (-0.38 – 0.37) |
| Male CORT x Breeding stage | -0.15 (-0.85 – 0.56) | **0.74 (0.12 – 1.36)** |
| Female CORT x Breeding stage | 0.18 (-0.48 – 0.85) | 0.36 (-0.29 – 1) |
| Male CORT x Female CORT x Breeding stage | 0.11 (-0.42 – 0.65) | -0.23 (-0.76 – 0.3) |

<sup>†</sup> *Estimates based on standardized data*
\* *Difference between incubation stage and nestling care*
\*\* *Difference from adult – to – yearling*

b. Number of fledglings
|  | *Model with baseline CORT* | *Model with stress-induced CORT* |
| --- | --- | --- |
| Fixed effects | *estimates<sup>†</sup> (95% CrI)* | *estimates<sup>†</sup> (95% CrI)* |
| Intercept | 3.47 (2.49 – 4.46) | 4.05 (3.18 – 4.92) |
| Male CORT | 0.12 (-0.48 – 0.71) | **-0.70 (-1.22 – -0.19)** |
| Female CORT | -0.07 (-0.73 – 0.59) | -0.08 (-0.65 – 0.49) |
| Breeding stage* | 0.04 (-1.09 – 1.18) | -0.33 (-1.33 – 0.68) |
| Laying date | **-4.13 (-6.66 – -1.6)** | -1.21 (-3.65 – 1.22) |
| Laying date<sup>2</sup> | **4.22 (1.76 – 6.69)** | 1.16 (-1.23 – 3.558) |
| Female condition | **-0.66 (-1.13 – -0.18)** | -0.27 (-0.69 – 0.16) |
| Male condition | 0.34 (-0.11 – 0.8) | -0.32 (-0.79 – 0.16) |
| Female age class** | -0.56 (-1.43 – 0.32) | -0.66 (-1.46 – 0.15) |
| Male age class** | 0.51 (-0.43 – 1.46) | 0.31 (-0.47 – 1.11) |
| Male CORT x Female CORT | **0.76 (0.23 – 1.3)** | 0.07 (-0.44 – 0.59) |
| Male CORT x Breeding stage | 0.12 (-0.72 – 0.96) | **1.11 (0.34 – 1.88)** |
| Female CORT x Breeding stage | 0.50 (-0.32 – 1.32) | **0.87 (0 – 1.74)** |
| Male CORT x Female CORT x Breeding stage | **-1.15 (-1.8 – -0.49)** | **-0.92 (-1.61 – -0.23)** |

<sup>†</sup> Estimates based on standardized data
\* Difference between incubation stage and nestling care
\*\* Difference from adult – to – yearling

**Figure 3.**
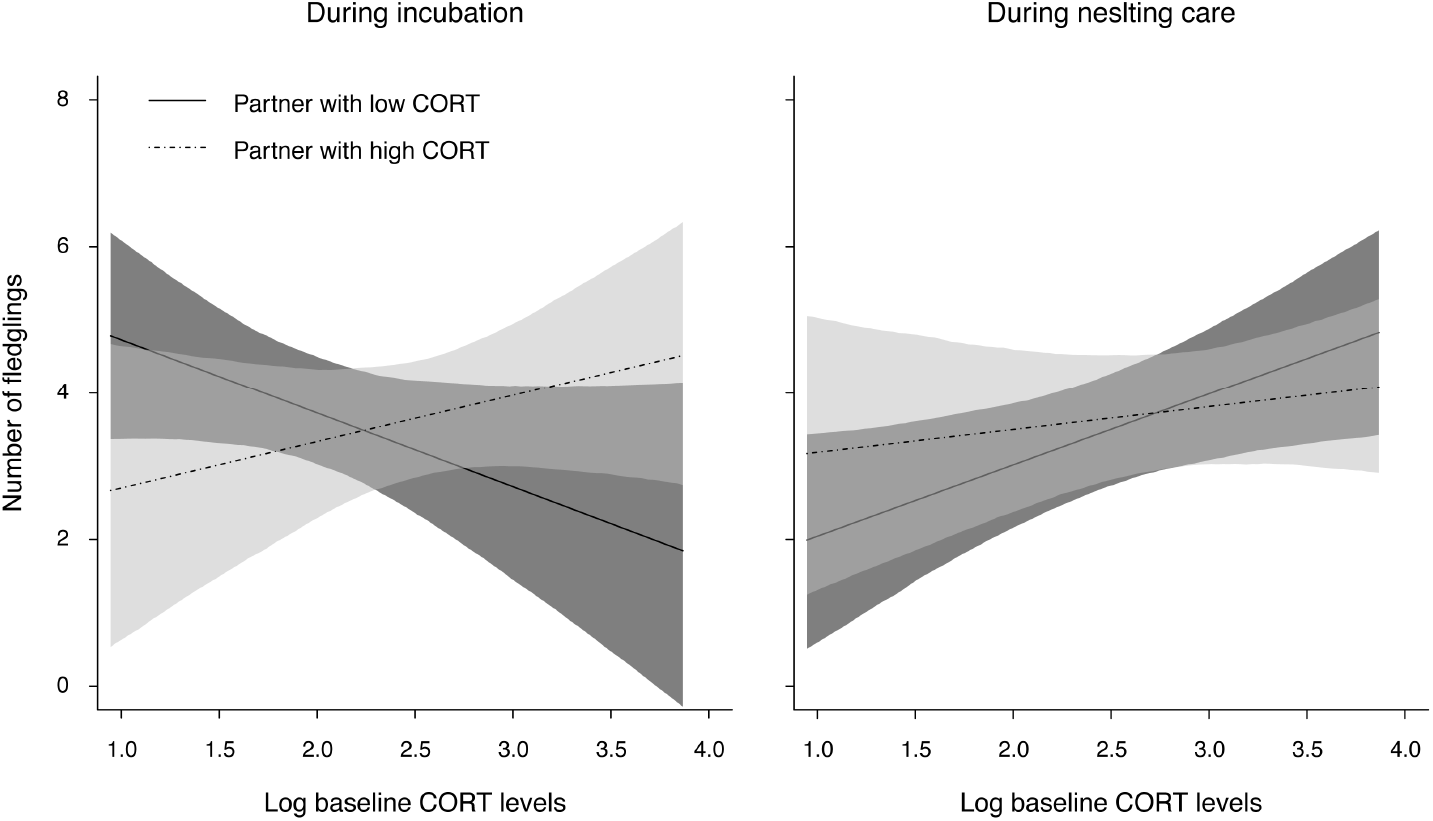
Relationship between the number of fledglings and log baseline CORT levels (ng/ml) of parents measured during incubation and nestling care. The black and the dotted lines represent the mean posterior distribution estimated from the baseline models (Table 2) for when the partner has low CORT levels (1^st^ quantile of males) and high CORT levels (3^rd^ quantile of males), respectively. The black and grey shadings represent the 95% CrI of the regression lines. For the figures, we considered the mean effect of laying date, age and condition of parents on number of fledglings.

**Figure 4.**
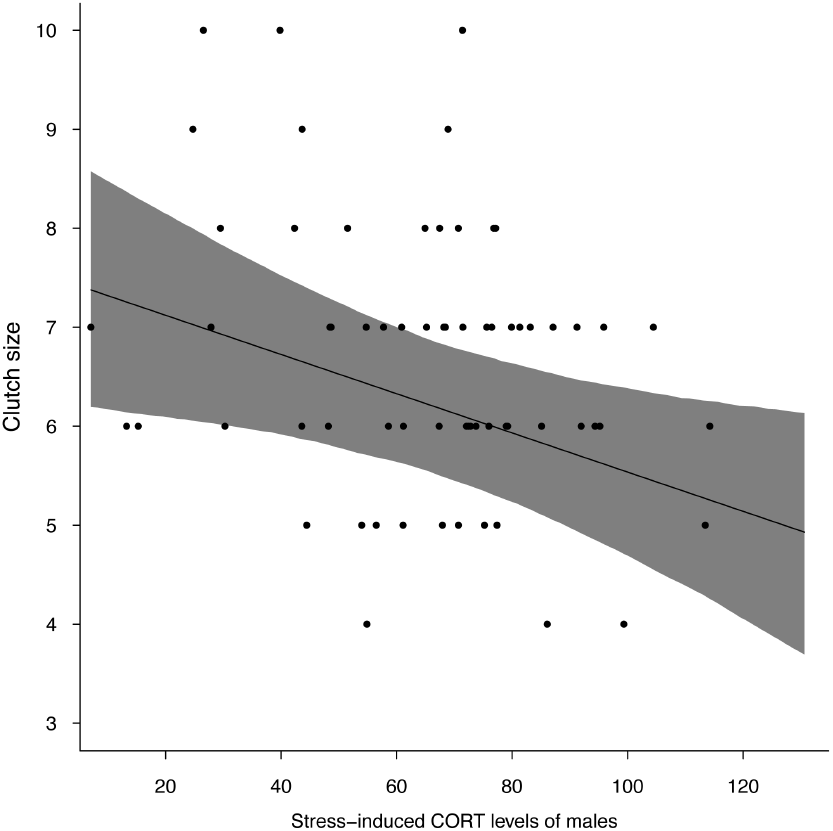
Clutch size in relation to stress-induced CORT response (ng/ml) of males during incubation. The black line represents the mean posterior distribution estimated from the stress-induced models (Table 2a) for when female partner has average CORT levels. The black shading represents the 95% CrI of the regression lines. For the figure, we considered the mean effect of laying date, age and condition of parents on clutch size.

**Figure 5.**
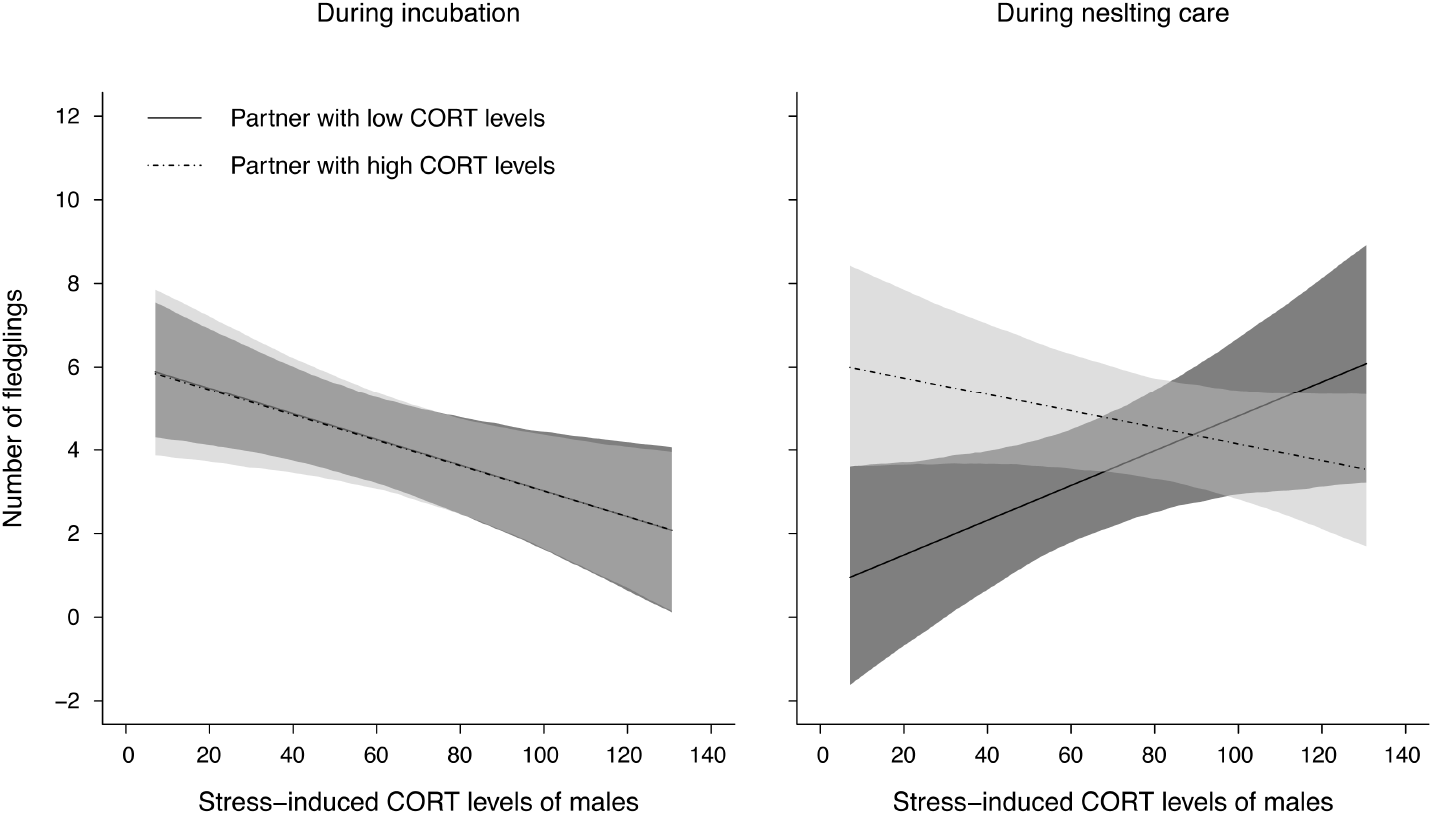
Number of fledglings in relation to stress-induced CORT response (ng/ml) of males measured during incubation and nestling care. The black and the dotted lines represent the mean posterior distribution estimated from the stress-induced models (Table 2) for when the partner has low CORT levels (1^st^ quantile of males) and high CORT levels (3^rd^ quantile of males), respectively. The black and grey shadings represent the 95% CrI of the regression lines. For the figures, we considered the mean effect of laying date, age and condition of parents on rearing success.

### Corticosterone in relation to nestling body mass

Variation in nestling body mass was associated with the maternal CORT levels measured during the incubation and nestling care in interaction with brood size but not with paternal CORT levels (Table 2) and was mostly observed in females presenting high CORT levels compared with mothers with low CORT levels. Thus, mothers with small broods and high baseline CORT levels had on average lighter nestlings than females with large broods and high baseline CORT levels, whereas nestlings raised by females with low baseline CORT levels had similar body mass regardless of the size of the brood (Figure 6). The stress-induced CORT response was similarly associated to the mass of nestlings as for baseline CORT levels (Table 2).

**Figure 6.**
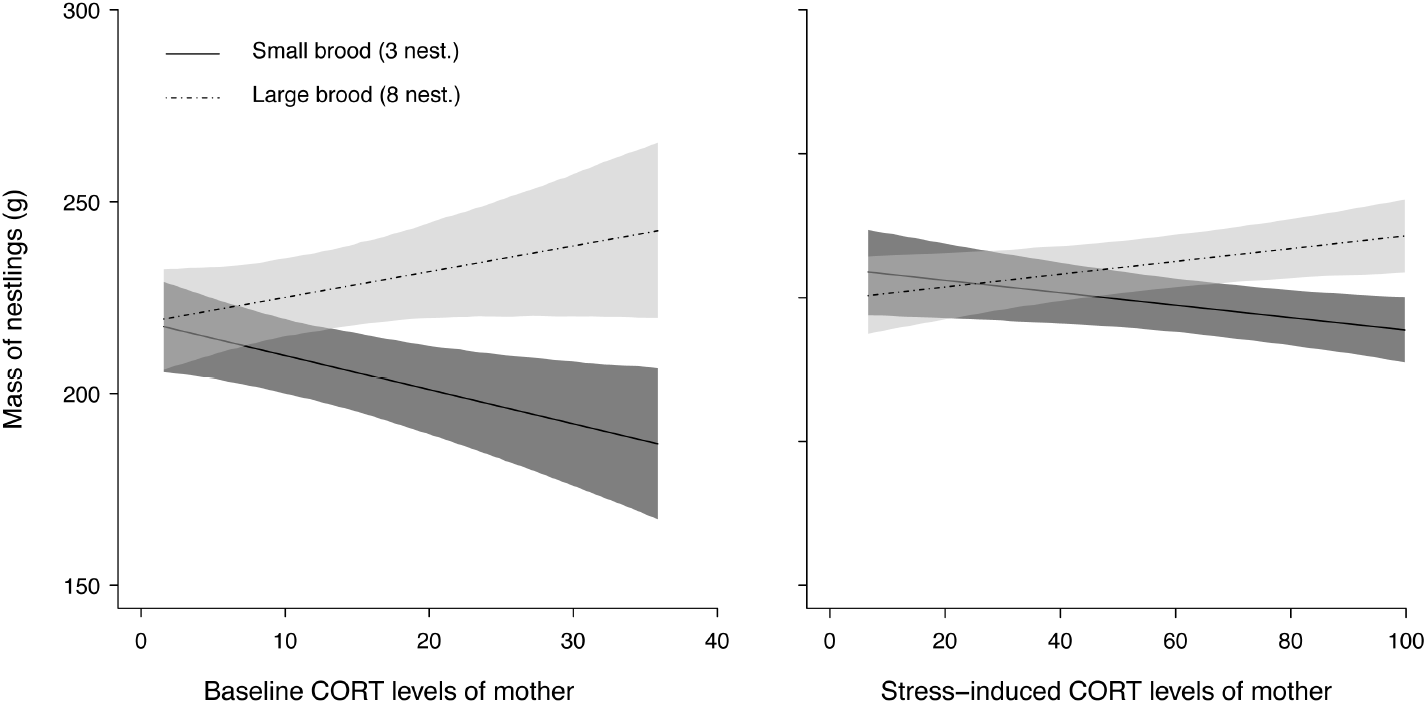
Relationship between body mass (g) of barn owl nestlings between age of 5 to 30 days with baseline and stress-induced CORT levels (ng/ml) of barn owl mother in interaction with size of the brood. The black and the dotted lines represent the mean posterior distribution estimated from baseline and stress-induced CORT models (Table 3) for small and large broods, 3 and 8 nestlings respectively. The black and grey shadings represent the 95% CrI of the regression lines. The regression lines and 95% CrI were drawn from 5’000 simulations of pairs of intercepts and slopes based on the posterior distribution of the model parameters. For both figures, we considered the mean effect of date, hour, sex age and rank of nestlings on body mass.

**Table 2.** Relationship between body mass of barn owl nestlings between aged 5 to 30 days and baseline and stress-induced CORT levels of male and female breeding partners. Effect of different parameters are estimated from a linear mixed effect model based on 715 and 721 body mass measurements (in 336 and 467 nestling barn owls from 74 and 95 broods) measured during 10 years between 2004 and 2016 for baseline and stress-induced CORT levels, respectively. Estimates (95% Bayesian credible intervals) of predictors are based on the draw of 50’000 random values from the joint posterior distribution of the model parameters. The year, identity of nestlings, parents and broods were added as random terms in the model to correct for repeated measurements. Predictors with a significant effect on mass of nestlings are written in bold. Nestlings were heavier at the beginning than at the end of the breeding season, first-born nestlings were heavier than last-born nestmates and nestlings raised in large broods were on average heavier than nestlings raised in smaller broods.

|  | <u>Model with baseline CORT</u><br><u>levels</u> | <u>Model with stress-induced</u><br><u>CORT levels</u> |
| --- | --- | --- |
| Fixed effects | *estimates<sup>†</sup> (95% BCrl)* | *estimates<sup>†</sup> (95% BCrl)* |
| Intercept | 217.76 (207.08 – 228.43) | 205.20 (196.84 – 213.57) |
| Date of sampling | -1.96 (-6.78 – 2.86) | -1.89 (-5.61 – 1.84) |
| Hour of sampling | **21.40 (6.79 – 36.19)** | **22.13 (8.97 – 35.39)** |
| Hour of sampling<sup>2</sup> | **-23.87 (-38.98 – -8.88)** | **-26.09 (-39.39 – -12.77)** |
| Age of nestlings | **76.24 (73.81 – 78.67)** | **73.00 (70.88 – 75.11)** |
| Sex of nestlings* | **10.05 (4.65 – 15.42)** | **9.82 (5.46 – 14.24)** |
| Rank of nestlings | **-9.62 (-12.37 – -6.88)** | **-9.70 (-12.06 – -7.36)** |
| Brood size | 5.65 (2.33 – 8.98) | 5.60 (2.57 – 8.59) |
| Mother CORT | -2.52 (-11.34 – 6.37) | 0.68 (-5.47 – 6.8) |
| Father CORT | 3.48 (-3.91 – 10.74) | 0.83 (-5.37 – 7.06) |
| Breeding stage** | -0.72 (-12.45 – 10.85) | -5.85 (-14 – 2.36) |
| Mother CORT x Brood size | **5.59 (1.65 – 9.51)** | **4.16 (1.46 – 6.86)** |
| Father CORT x Brood size | -3.51 (-7.81 – 0.82) | -2.10 (-5.26 – 1.1) |
| Father CORT x Mother CORT | -3.04 (-14.09 – 7.95) | 1.73 (-4.23 – 7.66) |
| Mother CORT x Breeding stage | 2.45 (-7.67 – 12.54) | -0.25 (-8.36 – 7.92) |
| Father CORT x Breeding stage | -3.49 (-13.64 – 6.72) | -1.36 (-9.61 – 7.15) |
| Father CORT x Mother CORT x Breeding stage | 0.49 (-11.39 – 12.43) | -2.36 (-10.19 – 5.47) |

<sup>†</sup> *Estimates based on standardized data*
\* *Difference from male – to – female*
\*\* *Difference between incubation stage and nestling care*

## Discussion

Our study shows that male and female partners tend to have similar CORT levels and that parental baseline and stress-induced CORT are associated with reproductive success, suggesting that parental CORT levels, or a correlated variable, mediate barn owl reproductive biology. However, CORT levels and reproductive parameters were related in a quite complex way, as it depended on when CORT was measured (during the incubation or nestling care) and the associations were different with male and female CORT levels (Figure 7).

**Figure 7.**
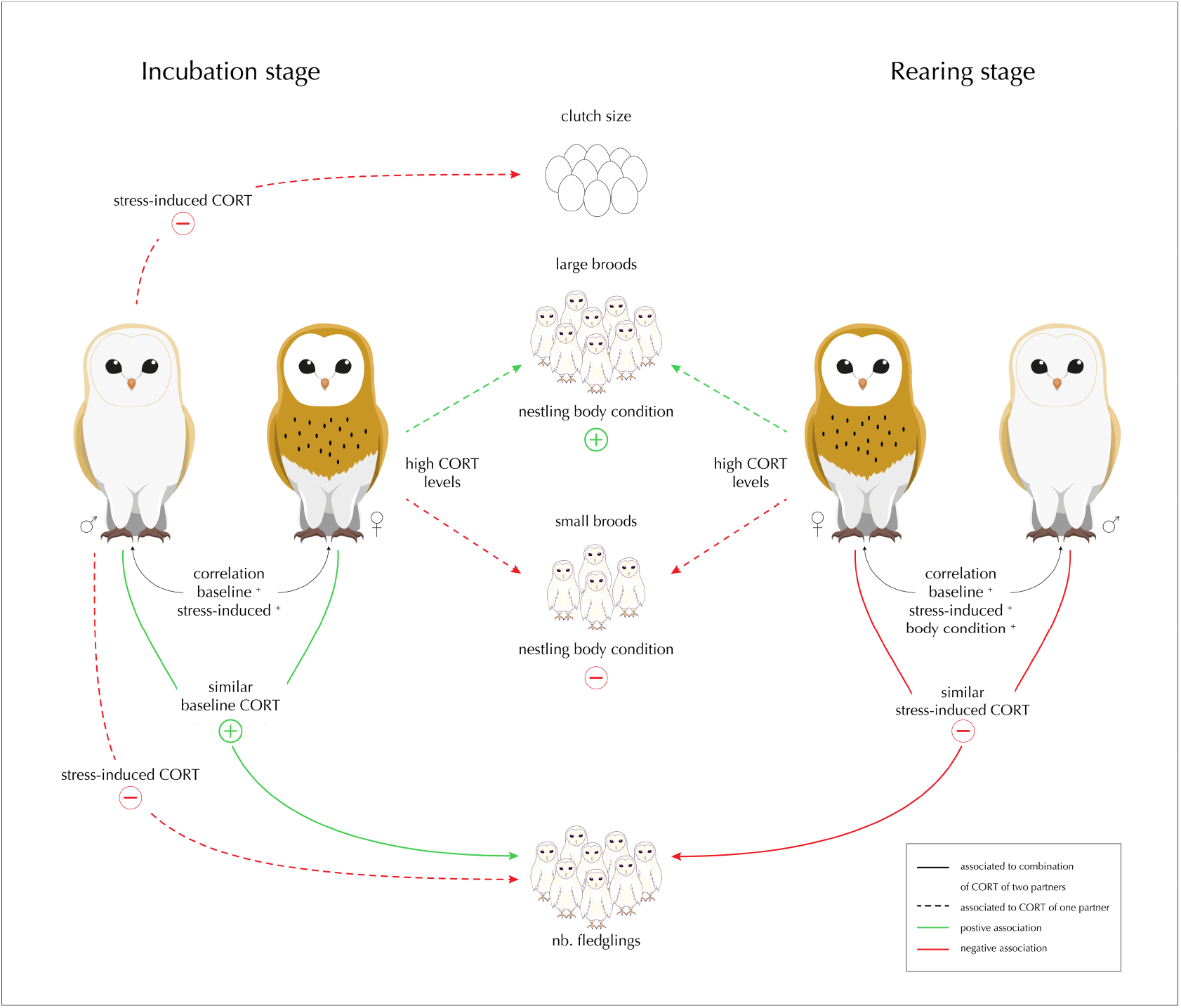
Summary of baseline and stress-induced CORT levels relationships with different fitness traits (*i.e*., clutch size, number of fledglings and offspring condition) in the barn owl. The plain arrows represent the effect associated to the combination of CORT levels of two partners and dashed arrows the effect associated to one of the partners only (Table 1). Green arrows represent positive relationships between CORT and fitness trait and red arrows negative relationships. The arrows between partners represent the correlation of CORT phenotypes and condition of partners (*i.e*., mass/wing). The (^+^) sign means that traits were positively and significantly correlated between partners. The association of CORT with the condition of nestlings (body mass) represents the interaction of female baseline and stress-induced CORT levels with brood size (Table 2).

### CORT levels and parental body condition

The positive correlation between CORT levels measured in male and female barn owls (Figure 1) can be explained by three non-mutually exclusive hypotheses. First, individuals may pair in an assortative way with respect to hormone profiles. Second, partners may have similar corticosterone levels because they exploit the same habitat and both take care of their progeny (*i.e*., raise the same number of nestlings). For instance, Ouyang et al. (2014) found in great tits that similarity in baseline CORT levels between pair members increased with the years of partnership and could potentially guarantee bond stability or, alternatively, could be the result from breeding in the same territory. Third, partners may mutually adjust their hormone profiles to each other to coordinate reproductive activities (Gabriel and Black 2012; Hirschenhauser et al. 1999) as shown in convict cichlid, *Amatitlania siquia*, in which breeding partners increase spawning size and their fitness by adjusting their behaviour (Laubu et al. 2016) and 11-ketestosterone levels to each other, a hormone which is involved in the gametogenesis and spawning behaviour of several fish species (Schweitzer et al. 2017). Because our results are correlative, we are not able to distinguish between these three hypotheses (assortative pairing, shared environment or post-pairing adjustment). These hypotheses could be further evaluated by sampling the same pairs at different stages of reproduction.

### Baseline corticosterone of parents and reproductive success

Whereas clutch size was not related to parental baseline CORT levels, parents produced more fledglings if they had similar baseline CORT levels during the incubation stage, a relationship that was not evident during the nestling stage. Baseline CORT is often associated with the degree of parental care (Bonier et al. 2011; Ouyang et al. 2013b) and nestling growth (Bonier et al. 2009b) and thereby, thought to influence reproductive success. From our results, we suggest that the number of fledglings is related to the degree to which parents coordinate their investment in parental care prior to the hatching period but not to hatching success. Indeed, hatching rate is not a key factor in fledgling success compared to nestling mortality during the provisioning period (*i.e*., 85% of pairs have a 100% hatching success and only 3% have less than 80% of hatching success). Partners having similar CORT levels may invest similar effort to raise their offspring, which could reduce the intensity of conflicts compared to partners that do not have alike CORT levels. Partners with dissimilar baseline CORT levels may indeed achieve a low reproductive success because pair members might not equally invest in parental care. Adults might thus pair preferentially with partners that are behaviourally or/and hormonally alike (Both et al. 2005; Ouyang et al. 2014; Schuett et al. 2011b). As shown by our results, reproductive success may not depend on the absolute baseline CORT levels (*i.e*., high or low baseline CORT) of pairs members as long as they are similar (Figure 3). Indeed, pairs with low or high baseline CORT levels may achieve similar reproductive success since they adjust their baseline CORT levels to the prevailing environmental condition, quality of their habitat or condition. For instance, pairs living in high quality habitat may have low baseline CORT levels and high fitness, whereas high quality pairs living in less favourable habitats may still be able to increase reproductive effort and baseline CORT levels to achieve similar reproductive success. Another possibility is that high quality pairs can raise large broods and increase baseline CORT to meet the increased energetical demand associated to their reproductive tasks.

### Stress-induced corticosterone of parents and reproductive success

Males showing lower stress-induced CORT responses during the incubation stage have larger clutches, and produce more fledglings than males with a higher stress response (Figure 5). These results might be explained by a potential link between stress response levels and habitat quality. For instance, barn owl nestlings living in intensive farming areas show an increased release in corticosterone after a stress event (Almasi et al. 2015), an effect that may also prevail in adult barn owls. Alternatively, mounting a high CORT stress response may be incompatible with a high investment in reproduction. Indeed, being highly sensitive to environmental challenges may divert males from their reproductive duties (Almasi et al. 2008). For instance, individuals that are highly sensitive to predation risk may be more prone to abandon their clutch (Love et al. 2004; Spee et al. 2011; Wingfield and Sapolsky 2003). The lack of association between female response to stress during the incubation stage and reproductive success indicates that if a clutch is abandoned this might be because the male rather than the female deserts the nest if disturbed during incubation. Tenacity to incubate the eggs by the female (the male does not incubate the eggs in the barn owl) may have selected females to not be too responsive to any source of environmental stress and during incubation dealing with potential predators may be the role of the father rather than the mother. This could explain why females most often remain on the eggs when disturbed and why more experienced males and those in better condition tend to have larger clutches (Table 1). These speculations should be further evaluated by manipulating predation risk to investigate male and female reactions.

Once the offspring have hatched, reproductive success depends on the combination of stress responses deployed by the two partners. During the nestling care period, partners with dissimilar stress-induced responses achieved a higher reproductive success than pairs with similar stress responses, implying that one but not the two partners should be able to mount a strong stress-induced CORT response. Thus, one parent should strongly react during a sudden stressful event, for instance in the presence of a predator, whereas the other partner should continue to invest effort to feed and protect the offspring. By showing different stress-responses, male and female partners may ensure that at least one parent responds correctly to the wide repertoire of environmental challenges that can require different responses. For instance, stress responses that are adapted to one situation, like predator attacks, may not be appropriate to other situations, such as inclement weather, food shortage or social strife (Coleman and Wilson 1998). Given that the stress-induced CORT response can regulate different behavioural traits and be modulated in relation to different life history stages (Romero and Wingfield 2016), it appears that the CORT response and personality traits of an individual are often linked (e.g., Baugh et al. 2012; Carere et al. 2003; Furtbauer et al. 2015; Pottinger and Carrick 2001; Stowe et al. 2010). For instance, individuals that are highly sensitive to stress by mounting a strong CORT response to a stressful challenge are commonly slow explorers and shy in social interactions and in front of predators. In contrast, individuals that are more resistant to stress (*i.e*., that mount a weak CORT response to a stressful challenge) are fast explorer and bold. Therefore, by pairing with a partner having a different sensitivity to stress, a pair may be composed of a resistant territory defender (*i.e*., fast explorer - bold) and a wiser decision-maker and efficient forager (*i.e*., slow explorer - shy) (Chittka et al. 2003; Moiron et al. 2016; Sih and Del Giudice 2012; Tan et al. 2018), who copes better with environmental changes (Cockrem 2013). They may also get the benefit of having a partner with a different and complementary foraging territory (Patrick and Weimerskirch 2014; see also Wolf and Weissing 2012). Thus, if with respect to baseline CORT levels partners should be similar or similarly adjust their baseline CORT levels, partners should have opposite patterns of stress responses to ensure a high reproductive success.

### Corticosterone of parents and nestling body mass

Previous study in barn owls showed that nestlings raised in large broods are usually heavier than nestlings raised in small broods (Almasi and Roulin 2015). This result suggests that when resources are abundant breeding pairs invest in large clutches and have nestlings in good condition. However, reproductive investment (e.g. food provisioning) not only depends on the environmental conditions, quality or availability of resources in the habitat but also on the capacity of parents to deal with the challenges of reproduction (Schifferli et al. 2014). Nestling provisioning is energetically demanding and hence, only high-quality parents might be able to mobilize enough resources to support the cost associated to the effort to raise large broods. Parents must also be able to adjust to fluctuation in environmental conditions, including stressful challenges (e.g., predator attack, climatic conditions, food availability), in a way that does not jeopardize reproductive success and their own survival. Therefore, parents should be able to adjust reproductive investment according to the prevailing environmental conditions, the energetic demand of the brood and their capacity to respond to a stressful challenge to maximizing their fitness.

Baseline CORT is involved in the mobilization of energetic resources of an organism (Sapolsky et al. 2000) and in several bird species, an elevated baseline CORT level is associated with an increased foraging behaviour and parental effort in different bird species (e.g., Angelier et al. 2008; Angelier et al. 2007; Bonier et al. 2011; Crossin et al. 2012). However, under certain circumstances like in poor-quality habitats CORT levels can increase and stay high (e.g., Kitaysky et al. 2007; Marra and Holberton 1998; see also Romero and Wingfield 2016; Scheuerlein et al. 2001), which can potentially affect the health (Bremner 2007; de Kloet et al. 1999; Martin 2009; Sapolsky et al. 2000) and reproductive success of an individual (Bonier et al. 2009a; Breuner et al. 2008). For instance, an experimental increase in baseline CORT can reduce the rate at which parents provision their offspring with food (Almasi et al. 2008) and ultimately the number of nestlings that fledge successfully (Silverin 1986). In our study, the effect of parental baseline CORT on offspring body mass varied depending on brood size. Mothers with low CORT levels and small broods had nestlings of similar weight as females with low CORT levels but large broods. In contrast, mothers with larger broods (*e.g*., 8 nestlings) than average (5 nestlings) and high CORT levels produced nestlings in better condition than females with smaller broods (*e.g*., 3 nestlings) and high CORT levels, an effect that was stronger when CORT levels were measured during nesting care than during late incubation stage. This pattern might be explained by the condition of the mother or quality of the habitat in which parents breed and their capacity to invest in reproduction. Mothers with large broods and high baseline CORT levels might have a higher capacity to support reproductive challenges (*i.e*., CORT-adaptation hypothesis, Bonier 2009a), as they may be in better condition or have access to more resources to support the cost of reproductive effort and, therefore have heavier nestlings. In contrast, females with small broods and high CORT levels live probably in a poor habitat and might be in poorer condition and, therefore, are not able to invest so much resources in reproduction (Almasi et al. 2013) and consequently have lighter offspring. This is in line with the finding that females in good condition had lower baseline CORT levels during incubation and higher baseline CORT levels during nestlings provisioning compared to females in poorer condition (Table S1). Surprisingly, offspring body condition was not or only weakly associated with paternal baseline CORT levels. Potentially, this is because mothers rather than fathers are in charge of cutting preys and feeding offspring during their first weeks of life.

Contrarily to baseline CORT that regulates daily physiological processes, the CORT stress response is thought to facilitate an individual escape from challenging or life-threatening events such as inclement weather, predator attacks or food shortage (Wingfield et al. 1998). The CORT stress response is presumed to favour self-maintenance over reproductive investment and, hence individuals with a higher stress sensitivity reproducing in an unpredictable environment should be more prone to redirect resources to self-maintenance at the expense of reproduction than less sensitive individuals. Several studies have indeed shown that parents with lower stress-induced response provision nestlings at a higher rate and fledge more offspring than individuals with higher stress-induced CORT responses (Angelier et al. 2009; Love et al. 2004; Ouyang et al. 2011; Schmid et al. 2013; Vitousek et al. 2014). However, in our case, females having large broods and a stronger stress-induced response had nestlings in better condition than females having a lower stress-induced CORT response, a relation that was in the opposite direction for females with small broods (Figure 6). As for baseline CORT, our results may reflect different coping strategies in relation to the context (*e.g*., stress resistant vs. stress sensitive), with some individuals being able to face the cost of reproduction at the expense of self-maintenance, while others may favour self-maintenance at the expense of offspring quality. For instance, a small artificial CORT increase in male barn owls during nestling provisioning showed that individuals are not all equivalently sensitive to elevated CORT (Almasi et al. 2008). Melanic males are more stress resistant than less melanic individuals. These differences might also be associated to the personality of individuals, a trait that is often linked to the CORT response of an individual to a stressor, and which are thought to perform differently depending on the environmental conditions (Cockrem 2007; Dingemanse et al. 2004; Ellis et al. 2006). For instance, in great tits, survival of nestlings has been shown to fluctuate with personality of mothers and winter condition. In harsh winter (i.e., low food), mothers with intermediate personality had more recruits the following years than females with extreme personality (fast and slow explorers). An association that was in the opposition direction during good winters, with mothers having extreme personality having more recruits the following year (Dingemanse et al. 2004). In our case, females with larger stress-induced CORT response might be more successful during good years or when environmental conditions are favourable (as reflected by brood size) and therefore invest more in nestling care than when conditions are less advantageous. In contrast, individuals with low stress-induced CORT response may be more resilient to environmental conditions which may affect less parents’ behaviour and thereby offspring condition. The choice of investing into reproduction may also depend on the value of the brood (Lendvai et al. 2007). Large broods have a higher residual reproductive value, which may induce mothers to sacrifice self-maintenance for large compared to small broods.

## Conclusion

Most studies investigating the CORT-fitness relationship in biparental species usually considered each partner individually and not the interaction of paternal and maternal CORT. Our results emphasize the necessity to measure and consider whether the hormonal profiles of male and female partners are coordinated, how and whether this is linked to reproductive biology. An open question is whether pairing with respect to hormone profiles is random and how these mate choice decisions affect reproductive behaviour.

## Acknowledgements

We are grateful to all the field assistants who helped us collecting the data in the field. We are also thankful to the Swiss National Science Foundation who financed this research (grant n° grant n° 3100A0-104134 and 31003A-127057 to L. J. and n° 31003A-173178 to A. R.). Capture and blood samples were taken under the legal authorization of the ‘Service vétérinaire du canton de Vaud.

---

**Table S1.** Baseline corticosterone levels and stress-induced corticosterone levels in relation to different covariates in adult barn owls. Effects of different parameters are estimated from a linear mixed effect model based on 117 baseline and stress-induced CORT samples measured during 9 years between 2006 and 2016 for baseline and stress-induced CORT levels. Baseline CORT was log transformed to obtain a normal distribution of the data. Estimates (95% Bayesian credible intervals) of predictors are based on the draw of 50’000 random values from the joint posterior distribution of the model parameters. The year, individual and brood identity were added in the models as random terms to correct for pseudo-replication. Predictors with a significant effect on fitness components are written in bold.

|  | *<u>Baseline CORT model</u>* | *<u>Stress-induced CORT model</u>* |
| --- | --- | --- |
| Fixed effects | *estimates<sup>†</sup> (95% CrI)* | *estimates<sup>†</sup> (95% CrI)* |
| Intercept | 2.66 (2.14 – 3.14) | 59.32 (40.44 – 76.95) |
| Date of sampling | **-0.35 (-0.55 – -0.14)** | -6.44 (-12.64 – 0.13) |
| Hour of sampling | **0.22 (0.03 – 0.38)** | 1.86 (-4.03 – 7.3) |
| Hour<sup>2</sup> of sampling | **-0.42 (-0.64 – -0.18)** | -11.06 (-18.41 – -3.34) |
| Sampling time | **0.13 (0.02 – 0.23)** | 0.15 (-4.07 – 4.5) |
| Age class* | -0.17 (-0.42 – 0.09) | 2.05 (-6.61 – 10.69) |
| Body condition | 0.3 (-0.23 – 0.84) | **-18.21 (-35.37 – -0.88)** |
| Breeding stage** | -0.59 (-1.42 – 0.28) | 7.03 (-19.15 – 33.9) |
| Sex *** | 0.1 (-0.44 – 0.65) | 10.08 (-7.17 – 27.07) |
| Body condition x Breeding stage | **-1.29 (-2.28 – -0.32)** | -11.34 (-44.12 – 20.07) |
| Body condition x Sex | **-0.67 (-1.28 – -0.06)** | 9.87 (-10.84 – 30.54) |
| Breeding stage x Sex | 0.38 (-0.58 – 1.34) | 0.69 (-30.77 – 31.55) |
| Body condition x Breeding stage x Sex | **1.92 (0.88 – 2.96)** | 13.67 (-22.46 – 49.89) |

<sup>†</sup> Estimates based on standardized data
\* Difference from yearling – to – adult
\*\* Difference from female – to – male
\*\*\* Difference between incubation stage and nestling care

